# Criterial Learning and Feedback Delay: Insights from Computational Models and Behavioral Experiments

**DOI:** 10.1101/2024.11.16.623975

**Authors:** Matthew J. Crossley, Benjamin O. Pelzer, F. Gregory Ashby

**Affiliations:** School of Psychological Sciences, Macquarie University, Sydney, Australia; Macquarie University Performance and Expertise Research Centre, Macquarie University, Sydney, Australia; Independent Researcher; Department of Psychological & Brain Sciences, University of California, Santa Barbara

**Keywords:** response criterion, criterial learning, associative learning, categorization, procedural learning

## Abstract

The notion of a response criterion is ubiquitous in psychology, yet its cognitive and neural underpinnings remain poorly understood. To address this shortcoming, three computational models that capture different hypotheses about criterial learning were developed and tested. The time-dependent drift model assumes the criterion is stored in working memory and that its value drifts over time. The delay-sensitive learning model assumes that the magnitude of criterial learning is temporally discounted by feedback delay. The reinforcement-learning model assumes that criterial learning emerges from stimulus-response association learning without an explicit representation of the criterion, with learning rate also temporally discounted by feedback delay. The performance of these models was investigated under varying feedback delay and intertrial interval (ITI) durations. The time-dependent drift model predicted that long ITIs and feedback delays both impair criterial learning. In contrast, the delay-sensitive and reinforcement-learning models predicted impairments only with feedback delays. Two behavioral experiments, which tested these predictions, showed that human criterial learning is impaired by delayed feedback but not by long ITIs. These results support the delay-sensitive and reinforcement-learning models, and suggest that even in tasks that appear to rely on explicit, rule-based reasoning, criterial learning may have strong associative underpinnings.

## Introduction

The notion of a response criterion is ubiquitous in psychology. It is a key component of almost all decision models. For example, the hypothesis that even YES-NO detection decisions are determined by comparing the sensory magnitude to a response criterion that is under the observer’s control, rather than to a fixed absolute threshold, allowed signal detection theory to supplant classical threshold theory as the dominant model in psychophysics (Green & Swets, 1966). All models that include a response criterion assume its value is learned and can shift if changes are made to instructions or payoffs. So criterial learning is a fundamental component of almost all decision-making models. Despite its importance, however, the cognitive and neural mechanisms that underlie criterial learning remain poorly understood.

This article addresses this shortcoming through a combination of computational modeling and empirical data collection. Specifically, we develop and test three different computational models that make qualitatively different assumptions about how the criterion is learned. The models differ in the role they assign to working memory and in whether they treat the response criterion as a fundamental psychological construct, or instead assume that behavior is driven purely by stimulus-response associations without any criterion guiding responses. These models are then tested in two behavioral experiments. The modeling and empirical focus are on how feedback delays and the length of the intertrial interval (ITI) affect criterial learning. All criterial-learning models assume that updating (i.e., learning) of the criterion occurs during the time interval between feedback presentation and the stimulus presentation that defines the onset of the next trial. So feedback delay and the length of the ITI are the independent variables that most clearly differentiate the conflicting predictions of criterial-learning models. Furthermore, feedback delays are known to impair some forms of learning (i.e., procedural) much more than others (i.e., learning that relies on declarative memory), so feedback-delay manipulations offer a powerful method of disambiguating the nature of criterial learning (Ell et al., 2009; Maddox & Ing, 2005; Maddox et al., 2003).

Although the models could be tested in any task that depends on criterial learning, the two experiments we describe used a one-dimensional category-learning task. In such tasks, stimuli vary across trials on two or more dimensions – one relevant to categorization and one or more that are irrelevant. The observer’s goal is to identify the relevant dimension and learn the response criterion that maximizes accuracy. This task has been used in hundreds of studies, and all current models of performance in this task emphasize the role of criterial learning.

In the first behavioral experiment, participants were explicitly instructed about the relevant dimension, thereby isolating criterial learning from rule selection and switching processes. The results showed that short feedback delays impaired learning compared to immediate feedback. The second behavioural experiment used stimuli composed of binary features – where no criterial learning is needed – and did not instruct participants about the relevant dimension. Thus, this experiment isolated rule selection and switching from criterial learning. The results found that increasing feedback delay or ITI did not affect performance. These findings suggest that feedback delays impact criterial learning but not the discovery of the relevant stimulus dimension. More broadly, they indicate that criterial learning may recruit procedural learning, even in tasks that seem to rely on explicit, rulebased processes.

## Experiment 1: The Models

We developed three computational models, each representing different architectures of criterial learning, and examined their sensitivity to the duration of ITIs and feedback delay. Two of the models assume that the criterion is stored in memory and that response decisions are made by comparing the current stimulus-driven percept to this stored referent. Both of these models further posit that the internal representation of the criterion is updated following errors using a simple gradient-descent rule. The third model assumes that no criterion is stored nor used to generate responses. Instead, reinforcement learning is used to form stimulus-response associations and these associations drive responding without any appeal to the notion of a criterion.

For each of the three models, denote the time of stimulus presentation on the current trial by T_S_, the response time by T_R_, the time when feedback is displayed by T_F_, and the time when the stimulus that begins the next trial is displayed by 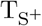. Then note that the feedback delay equals t_FD_ = T_F_ *−* T_R_ and the duration of the ITI equals 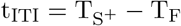.

### Time-Dependent Drift Model

The time-dependent drift model assumes that the observer constructs a criterion, holds this value in working memory, and then makes response decisions by comparing the percept to the criterion. The model also assumes that both the memory representation of the criterion and the perceived value of the stimulus gradually drift over time. The model further assumes that the extent of this drift increases with time, both within a trial and between consecutive trials.

Let *x*_*n*_(*t*) and *c*_*n*_(*t*) denote the values of the percept and the criterion, respectively, on trial *n* at time *t*. Then the decision rule on trial *n* is:

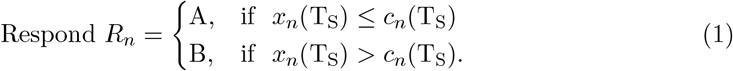

If positive feedback is received, then the value of the criterion remains unchanged for the next trial (except for drift – see Equation 3). If negative feedback is received, then the criterion is modified according to the standard model (Sutton & Barto, 1998):

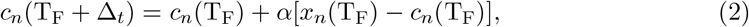

where *α* is a learning-rate parameter, and Δ_*t*_ is the time it takes to complete the updating. It is straightforward to show that the iterative Equation 2 is equivalent to computing a weighted mean (weighted by recency) of the values of all percepts that occur on error trials (e.g., Ashby, 2017). This updating rule will gradually converge on the optimal criterion value. Since this model assumes that criterial learning relies on working memory – and the available evidence suggests that logical reasoning and working memory are unaffected by feedback delays of several seconds (e.g., in one-dimensional rule-based category learning tasks; Ell et al., 2009; Maddox & Ing, 2005; Maddox et al., 2003) – we assume that the learning process described in Equation 2 is likewise unaffected by feedback delay.

Although the criterial-learning process described by Equation 2 is not affected by feedback delay, we assume that both the stimulus and the criterion representations drift randomly throughout the duration of time they are maintained in working memory. The representation of the criterion must always be maintained in working memory, whereas drift in the stimulus representation affects performance up until the feedback is presented, but not afterwards.

We modeled the drift in both the criterion and the percept by adding white noise to their initial values. Specifically, we assumed that for all *t >* T_S_

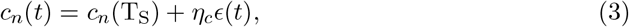

and

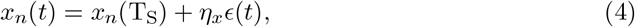

where *ϵ*(*t*) is white noise and *η*_*c*_ and *η*_*x*_ are parameters that determine the amount of drift over time. This model predicts that at the time of feedback, the response criterion *c*_*n*_(T_F_) will be normally distributed with mean *c*_*n*_(T_S_) and variance 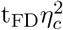. Similarly, at the time when the stimulus that defines the next trial is presented, the criterion 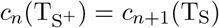 is normally distributed with mean *c*_*n*_(T_S_) and variance 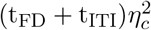. The predictions for the percept are similar. Specifically, at the time when feedback is presented, the percept *x*_*n*_(T_F_) is normally distributed with mean *x*_*n*_(T_S_) and variance 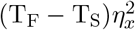.

### Delay-Sensitive Learning Model

Like the time-dependent drift model, the delay-sensitive learning model assumes that decisions are based on comparing the current stimulus to a stored referent. However, unlike the time-dependent drift model, the criterion remains stable over time and does not drift. For this reason, the memory system used to store the criterion may be different from working memory. In other words, *c*_*n*_(*t*) = *c*_*n*_ for all *t >* T_S_. This model also assumes that the magnitude of error-driven updates to the criterion decreases in proportion to the length of the feedback delay. As a result, when feedback is delayed, the system becomes less responsive to errors, leading to slower learning compared to immediate feedback conditions. This model also assumes that the percept does not drift over time. Instead, it models perceptual noise as a time-invariant perturbation. Specifically, the delay-sensitive learning model assumes the observer uses the following decision rule:

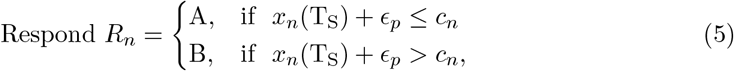

where the time-invariant perceptual noise term *ϵ*_*p*_ is normally distributed with mean 0 and variances 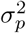. This same value of perceptual noise corrupts all future values of the percept. Therefore, *x*_*n*_(*t*) = *x*_*n*_(T_S_) + *ϵ*_*p*_ for all *t >* T_S_.

The delay-sensitive learning model assumes that longer feedback delays slow criteral learning. Specifically, the model assumes that if negative feedback is received, the criterion is updated as follows:

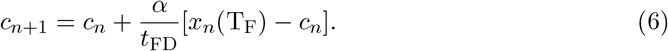

The scale factor (1*/t*_FD_) captures the notion that the length of the feedback delay slows the *rate* of procedural learning.

### Reinforcement-Learning Model

The reinforcement-learning model gradually associates responses with stimuli, and therefore does not include an explicit representation of the response criterion. The model is based on an actor-critic architecture and uses reinforcement learning to form stimulus-response associations (Sutton & Barto, 1998).

This model assumes that the perceptual representation of each stimulus is defined by the pattern of activation across 25 sensory units that are characterized by overlapping tuning curves. Each unit is maximally excited by one specific stimulus, which we call the unit’s preferred stimulus. Specifically, the activation of the *i*^th^ sensory unit on trial *n* is given by

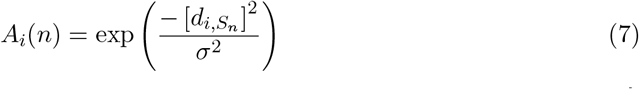

where 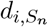 is the (Euclidean) distance between the preferred stimulus value of the *i*^th^ sensory unit and the value of the stimulus that was presented on trial *n*, and *σ* is a constant that increases with perceptual noise.

The model includes two decision or actor units – one associated with each of the two possible responses. Initially, each sensory unit is connected to both actor units with some random connection strength. Let *ω*_*iJ*_ (*n*) denote the strength of the connection between sensory unit *i* and actor unit *J* (for *J* = A or B) on trial *n*. Then the activation in actor unit *J* on trial *n*, denoted by *V*_*J*_ (*n*), equals

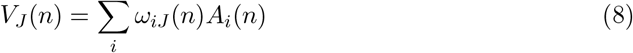

Responses are generated by the decision rule:

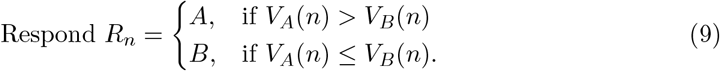

The connection strengths *ω*_*iA*_(*n*) and *ω*_*iB*_(*n*) are updated after feedback is received on each trial according to standard reinforcement learning rules (Sutton & Barto, 1998):

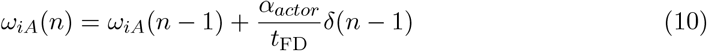

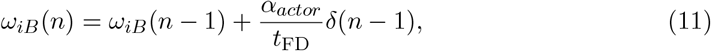

where *α*_*actor*_ is the learning-rate of the actor, and *δ*(*n −* 1) is the reward prediction error on trial *n −* 1. Note that as in the delay-sensitive learning model, the learning rate is scaled by the inverse of the feedback delay. Much evidence suggests that this type of stimulus-response learning is mediated largely within the striatum, and is facilitated by a dopamine (DA) mediated reinforcement learning signal that is time dependent (e.g., Valentin et al., 2014). Specifically, the dopamine signal generated by positive feedback appears to peak at around 500 ms after feedback and then decay back to baseline levels within 2 or 3 s (Yagishita et al., 2014). As a result, synaptic plasticity at cortical-striatal synapses is attenuated with increasing feedback delays (Yagishita et al., 2014). The scaling of *α*_*actor*_ by 1*/t*_FD_ models this phenomenon.

The reward prediction error on trial *n −* 1 is defined as the value of the obtained reward, denoted by *R*(*n−* 1) minus the value of the predicted reward, denoted by *P* (*n−* 1):

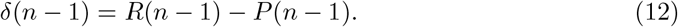

The predicted reward on trial *n* is determined by the critic via

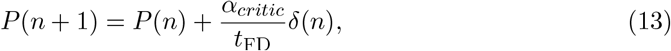

where *α*_*critic*_ is the learning rate of the critic. This learning rate is again scaled by the inverse of the feedback delay.

### Simulation Results

We investigated how changing the duration of the feedback delay and ITI affect criterial learning for each of these three models. More specifically, we simulated performance of each model in a categorization task that included two categories of stimuli that varied on a single stimulus dimension and that could be categorized perfectly by comparing each stimulus to an appropriate value of the response criterion (We used the same experimental design as in Experiment 1, so see those methods for more details). Our simulations included three experimental conditions: Delayed Feedback, Long ITI, and Control. The Delayed-Feedback condition included a long feedback delay (3.5 s) and a short ITI (0.5 s). The Long-ITI condition matched the total trial duration of the Delayed-Feedback condition but with a short feedback delay (0.5 s) and a long ITI (3.5 s). The Control condition included both a short feedback delay (0.5 s) and a short ITI (0.5 s). Each simulation continued for 200 trials or until the model responded correctly for 12 trials in a row. For each set of parameter values, we simulated the model 100 times and averaged the results, yielding one observed performance metric measured in trials-to-criterion for each condition. All scores were then normalized by the largest observed value, and as a result, all axes in the figures that describe the simulation results range from zero to one.

Our approach was to generate predictions for each of the three experimental conditions across a wide range of parameter values. We then used parameter-space partitioning (PSP) to evaluate the performance of each model (PSP; Pitt et al., 2006). PSP calculates the proportion of the parameter space where a model makes specific qualitative predictions. We focused on four such predictions. The first was that learning under feedback delay would be slower than in the other two conditions. The second was that learning in the Long ITI condition would be slower than in the other two. The third was that learning in the control condition would be slower than in either of the other two conditions. The fourth was a catch-all category encompassing any other possible patterns of results. For each model, the PSP analysis quantified the proportions of the explored parameter space where the model made these qualitative predictions.

### Time-dependent drift model

We investigated predictions of this model across a wide range of values for the parameters *η*_*x*_, *η*_*c*_, and *α*. Specifically, in the case of *α*, we stepped through every value in the interval [0, 1], with a step size of .01. In the case of both *η*_*x*_, and *η*_*c*_, we searched over the interval [0.1, 5], with a step size of 0.5. The results are shown in Figure 1.

**Figure 1.**
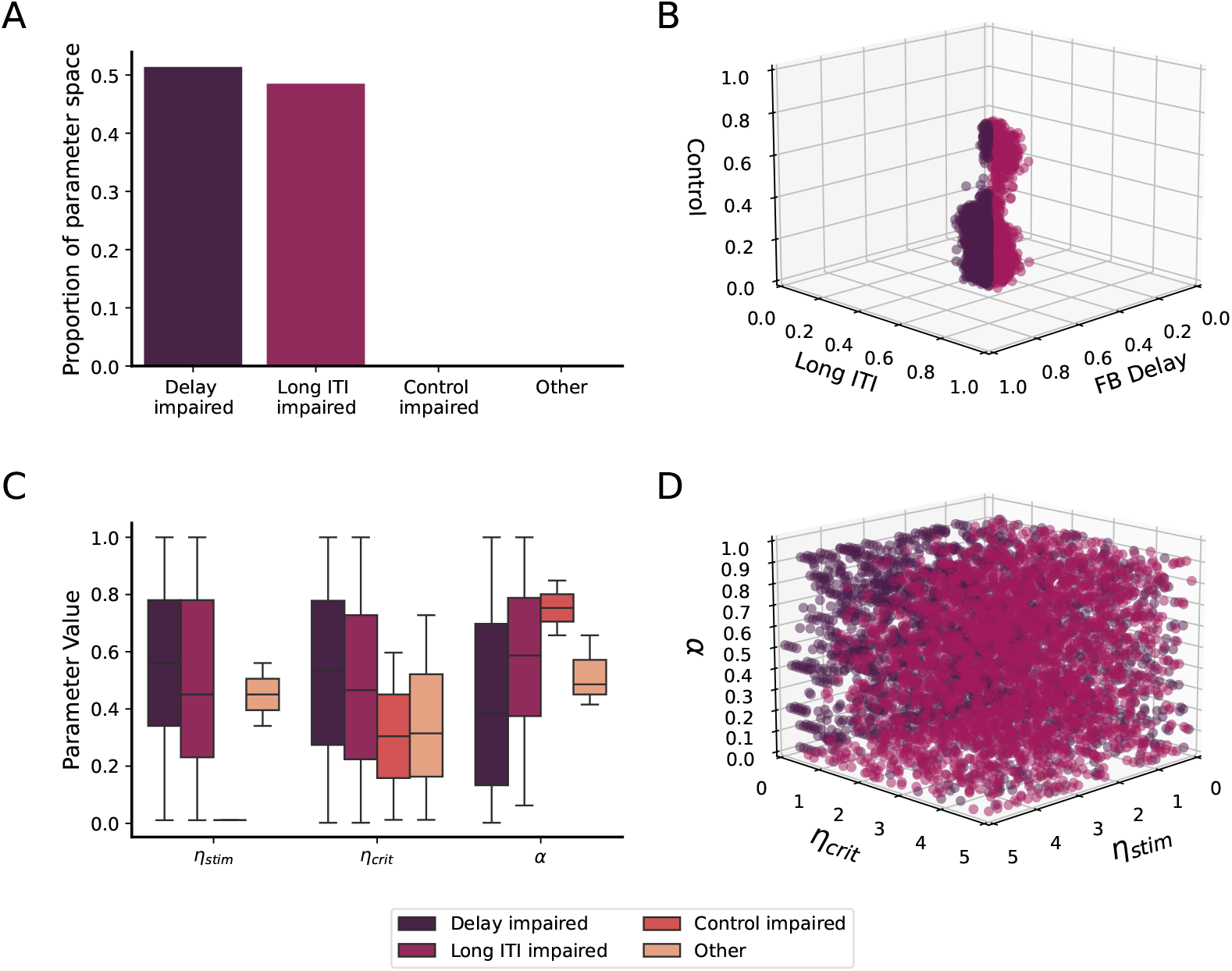
**A:** The proportion of parameter space where the time-dependent drift model predicted each of four qualitative patterns: (1) slower learning under feedback delay, (2) slower learning in the Long-ITI condition, (3) slower learning in the control condition, and (4) any other pattern. **B:** Simulated trials-to-criterion for each condition. All scores were normalized by the largest trials-to-criterion value within a given parameter set, so all axes range from zero to one. **C:** Boxplot of the parameter ranges leading to each PSP pattern. All parameter values were normalized by the largest value in the search range, so the ordinate ranges from zero to one for all parameters. **D:** Scatter plot of the parameter ranges associated with each PSP pattern. Note: In all panels, color indicates the PSP pattern.

Figure 1A shows that the model essentially always predicts that increasing the feedback delay or the ITI have similar detrimental effects on learning. The model makes this prediction because it assumes that the criterion drifts during both the feedback delay and during the ITI. So the critical variable for the model is the time between the response on trial *n* and the presentation of the stimulus on trial *n* + 1. The model assumes that the criterion drifts during this entire time, and therefore how this interval is divided between feedback delay and ITI is relatively unimportant to the model predictions.

### Delay-sensitive learning model

We simulated the performance of the delay-sensitive learning across a wide range of parameter values for *σ*_*p*_ and *α*. Specifically, in the case of *α* we stepped through every value in the interval [0, 1], with a step size of .1. In the case of *σ*_*p*_, we searched over the interval [0.1, 5], with a step size of 0.1.

The results are shown in Figure 2. Note that virtually all combinations of the parameter values predict that feedback delay impairs criterial learning more than increasing the ITI. This makes sense because Equation 6 shows that the delay-sensitive learning model predicts that increasing the feedback delay (i.e., increasing *t*_*F D*_) will impair learning – for any value of *α >* 0. Large values of *σ*_*p*_ will also impair performance because of distortion to the percept, but this interference will be the same in all conditions.

**Figure 2.**
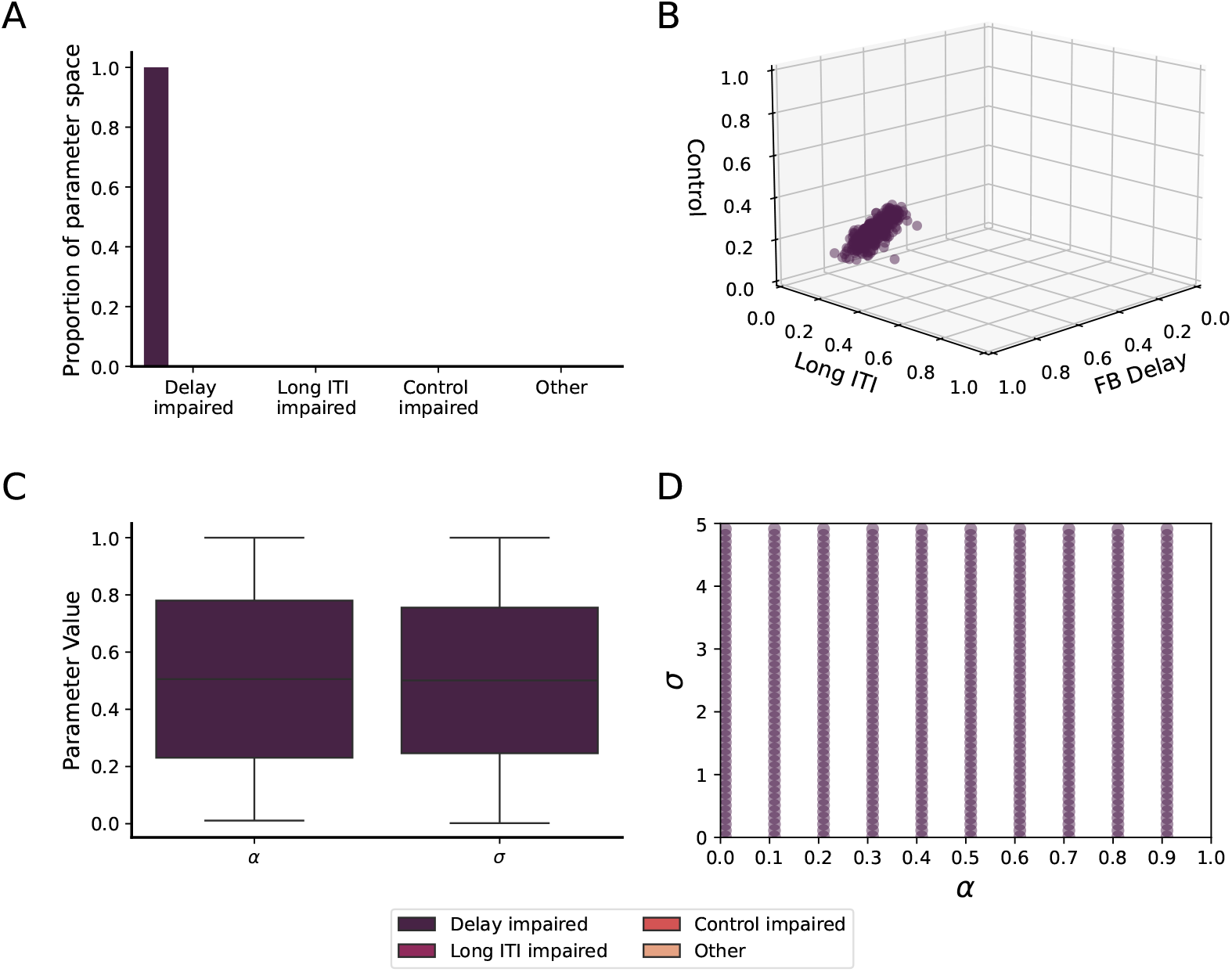
**A:** The proportion of parameter space where the delay-sensitive learning model predicted each of four qualitative patterns: (1) slower learning under feedback delay, (2) slower learning in the Long-ITI condition, (3) slower learning in the control condition, and (4) any other pattern. **B:** Simulated trials-to-criterion for each condition. All scores were normalized by the largest trials-to-criterion value within a given parameter set, so all axes range from zero to one. **C:** Boxplot of the parameter ranges leading to each PSP pattern. All parameter values were normalized by the largest value in the search range, so the ordinate ranges from zero to one for all parameters. **D:** Scatter plot of the parameter ranges associated with each PSP pattern. Note: In all panels, color indicates the PSP pattern.

### Reinforcement-learning model

We investigated the effects on performance predicted by the reinforcement-learning model of three parameters – the perceptual-noise variance *σ*^2^ and the actor and critic learning rates (i.e., *α*_*actor*_ and *α*_*critic*_, respectively). In the case of both *α*_*actor*_ and *α*_*critic*_, we stepped through every value in the interval [0, 0.2], with a step size of .01. We constrained our search over these parameters to this interval because reinforcement-learning models are prone to instability at very high learning rates (Sutton & Barto, 1998). As evidence of this, at higher learning rates, the model failed to learn with any consistency – that is, in most cases, it failed to reach the learning criterion (12 correct responses in a row) within the allowable 200 trials. In the case of *σ*, we searched over the interval [1, 10], with a step size of 1. As with the other models, all scores were normalized by the largest observed value.

The results are described in Figure 3. Note this model predicts that delayed feedback impairs criterial learning more than a long ITI across a wide volume of parameters settings. As can be seen in Figure 3C, the only exceptions tend to occur when the value of any of the three parameters approaches the maximum end of the range explored. The logic here is similar to the delay-sensitive learning model – that is, both models predict that increasing the feedback delay *t*_*F D*_ will impair learning – for all values of the learning rate.

**Figure 3.**
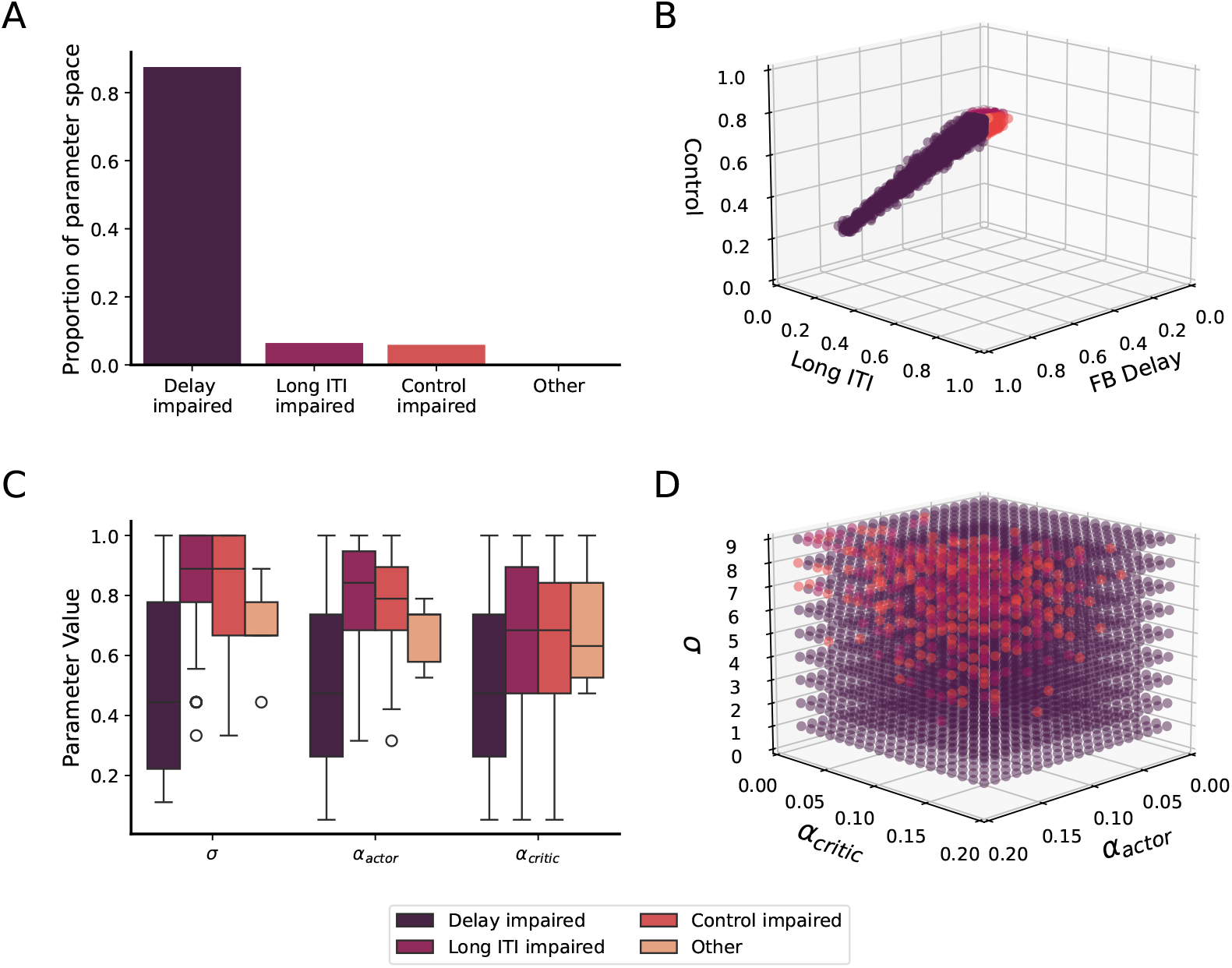
**A:** The proportion of parameter space where the reinforcement-learning model predicted each of four qualitative patterns: (1) slower learning under feedback delay, (2) slower learning in the Long-ITI condition, (3) slower learning in the control condition, and (4) any other pattern. **B:** Simulated trials-to-criterion for each condition. All scores were normalized by the largest trials-to-criterion value within a given parameter set, so all axes range from zero to one. **C:** Boxplot of the parameter ranges leading to each PSP pattern. All parameter values were normalized by the largest value in the search range, so the ordinate ranges from zero to one for all parameters. **D:** Scatter plot of the parameter ranges associated with each PSP pattern. Note: In all panels, color indicates the PSP pattern.

### Summary of modeling results

We investigated three criterial-learning models that make different predictions about how increases in the feedback delay and ITI affect learning. The time-dependent drift model predicts that the criterion always drifts, so the critical variable is the sum of the feedback delay and ITI. How this sum is divided into separate feedback delay and ITI time intervals has little effect on the model’s predictions. In contrast, the delay-sensitive learning model predicts that feedback delays are necessarily more detrimental to performance than increases in the ITI. Finally, the reinforcement-learning model also predicts that, in general, feedback delays should impair learning more than long ITIs, although this model is somewhat more flexible than the delay-sensitive learning model and can account for a small or null effect of feedback delay within a restricted region of its parameter space. The obvious next question is how human criterial learning is affected by these independent variables. Experiments 2 and 3 were designed to address this question, and therefore also to test the predictions of the three models.

## Experiment 2

Experiment 2 investigated how feedback delay and the length of the ITI affect criterial learning in humans. As a result, it also provides rigorous tests of the predictions of the time-dependent drift model, the delay-sensitive learning model, and the reinforcementlearning model.

The stimuli in Experiment 2 were circular sine-wave gratings that varied across trials in bar width and bar orientation. These stimuli were divided into two categories according to their value on one of the two dimensions. In other words, the optimal strategy was to set a response criterion on the single relevant dimension, and then choose a categorization response based on whether the value of the presented stimuli on this dimension was larger or smaller than the criterion value.

The experiment isolated criterial learning by (1) explicitly instructing participants on the relevant stimulus dimension and rule structure (e.g., thick bars are “A”, thin bars are “B”), and (2) eliminating variability along the irrelevant stimulus dimension. In other words, the instructions identified the relevant stimulus dimension, and as a result, the only learning required was criterial learning.

### Method

#### Apparatus

All experiments were performed in a dimly lit room. Participants sat approximately 24” from a 17” *×* 11” monitor running at a resolution of 1680 *×* 1050 pixels. Participants made category judgments by pressing the ‘d’ or ‘k’ keys on a standard computer keyboard for ‘A’ or ‘B’ choices, respectively. Stickers with bold print ‘A’ or ‘B’ were placed on the appropriate keys.

#### Stimuli and Categories

Stimuli were circular sine-wave gratings that varied in bar width and bar orientation, drawn from various 1-dimensional uniform distributions specific to the current category problem. We first defined an arbitrary 2-dimensional [0 *−* 100, 0 *−* 100] stimulus space, and then split each dimension of this space into 7 bins of width 14 units each. Each (*x, y*) pair from this arbitrary stimulus space was converted to a grating according to the nonlinear transformations defined by Treutwein et al. (1989), which roughly equate the salience of each dimension (for details, see also Crossley and Ashby (2015)).

The structure of the various criterial-learning tasks is illustrated in Figure 4. Each criterial-learning problem was created by first randomly selecting a relevant dimension, and then randomly selecting one of the 7 bins defined on that dimension. Each bin was also associated with a corresponding unique value on the irrelevant dimension. We buffered the to-be-learned response criterion by 10% of total bin width on either side with a no-stimulus region. Random uniform samples from the remaining eligible region of each bin were then selected and presented to the participant until 9 correct responses out of any 10 responses in a row advanced the participant to the next problem. Note that every category problem was a simple one-dimensional rule in which optimal accuracy was 100%. Note also that the relative location of the optimal response criterion varied across problems.

**Figure 4.**
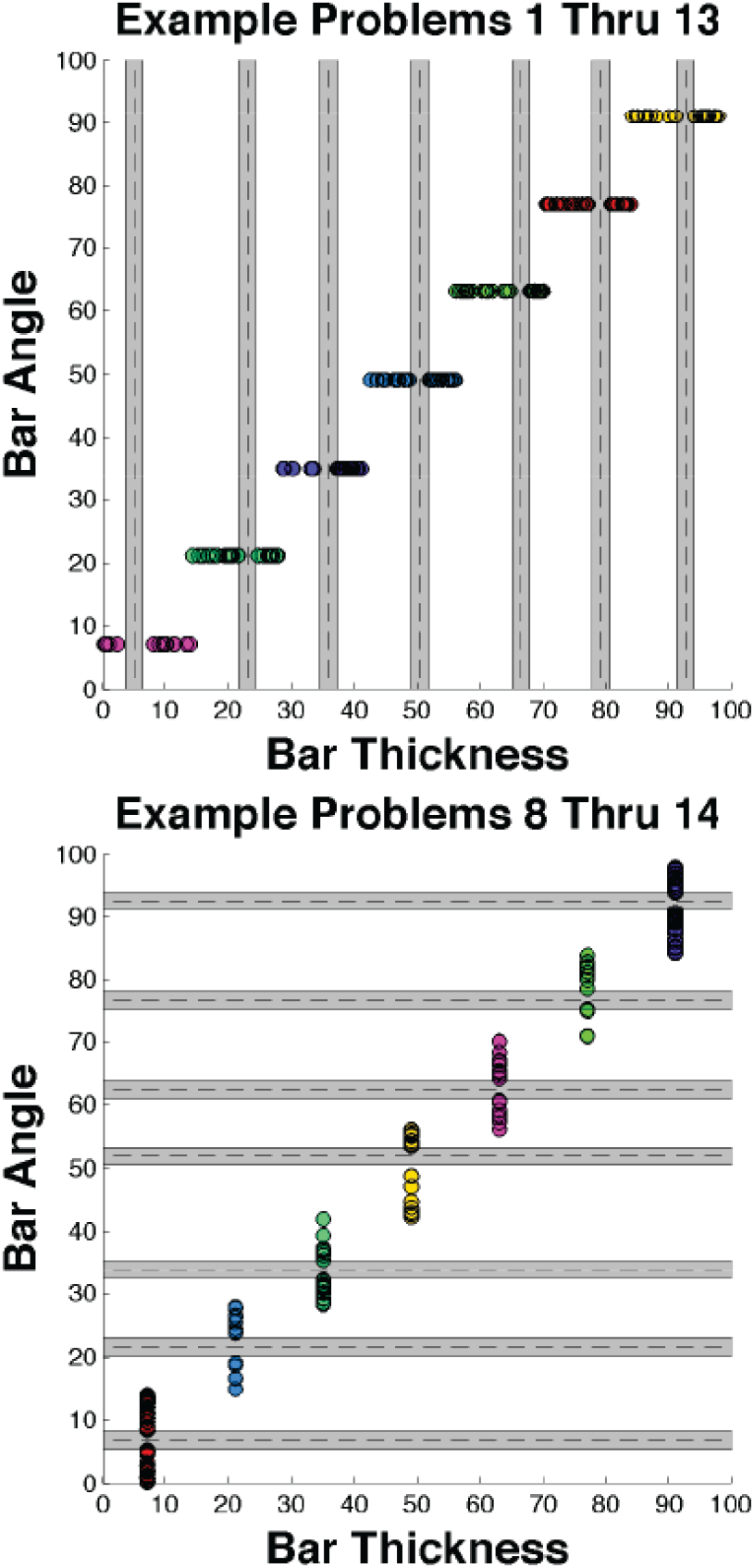
Category sample space. Different colors represent different category problems. Dashed lines are category boundaries (criterion) and the surrounding solid lines mark the no-stimulus region in which no stimuli were sampled.

#### Procedure

There were three conditions (described in detail in Table 1). In the Delayed-Feedback condition, feedback was delayed 3.5 s after the response and the ITI was 0.5 s. In the Long-ITI condition, feedback was delayed 0.5 s after the response and the ITI was 3.5 s. Finally, in the Control condition, feedback was delayed 0.5 s after the response and the ITI was 0.5 s.

**Table 1.** Durations (in s) of Trial Events in each Condition.

| Conditions | Stim | Mask | FB | ITI |
| --- | --- | --- | --- | --- |
| Control | RT | 0.5 | 1.0 | 0.5 |
| Delayed Feedback | RT | 3.5 | 1.0 | 0.5 |
| Long ITI | RT | 0.5 | 1.0 | 3.5 |

Each participant completed a series of one-dimensional category-learning tasks or problems, which are described in Figure 4. Each problem included stimuli in two distinct clusters that varied over a restricted range of the relevant dimension. Critically, the optimal criterion value varied from problem-to-problem with respect to its position within this range. For example, for some problems the optimal criterion was below the midpoint of the range and for other problems it was above the midpoint.

Participants were explicitly told the relevant dimension for each problem, as well as the generic response mapping (e.g., thick bars = “A”, thin bars = “B”). Figure 5 shows the structure of an example trial, along with an example of a typical category structure. All trials in every condition included a 500 ms fixation cross, a response-terminated stimulus, a circular white-noise mask, corrective feedback, and an inter-trial interval (ITI) that varied according to condition. The text ‘Correct’ was displayed in centered, large green font after correct responses, and the text ‘Incorrect’ was displayed in centered, large red font after incorrect responses.

**Figure 5.**
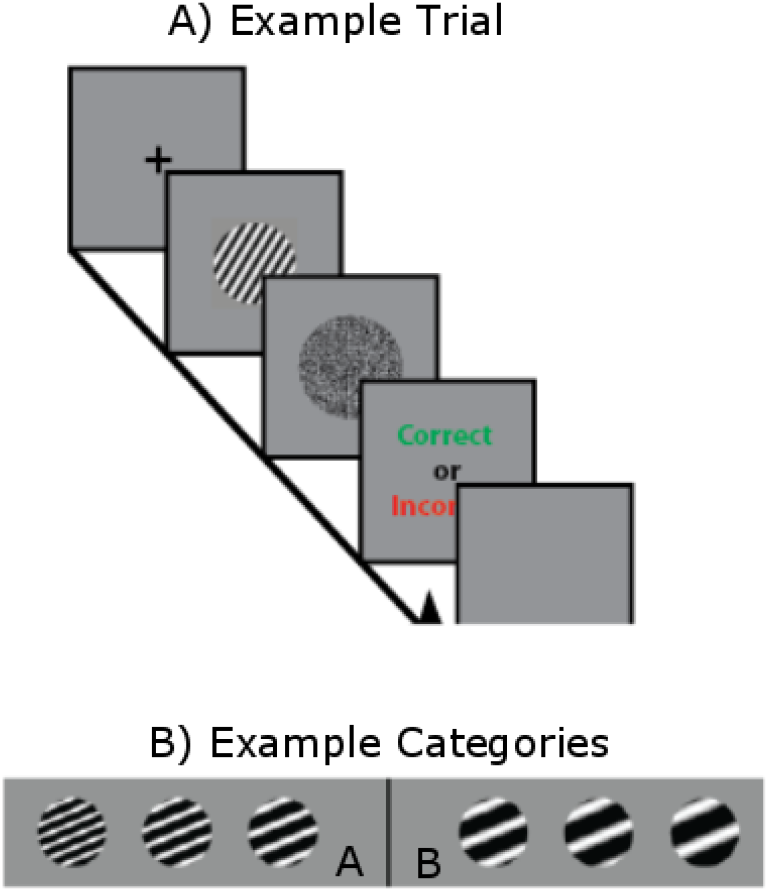
Example trial and category problem. A) Events that occurred on each trial. B) An example of a typical category structure.

Participants practiced each problem until they responded correctly on 9 of the previous 10 trials. At this point, the problem changed. Each participant completed as many problems as possible in 512 trials or until they had been in the lab for 60 minutes (including time to acquire consent and give instructions), at which point the session was terminated.

#### Participants

Fifty-nine participants participated in Experiment 2. All were UCSB undergraduates and received course credit for their participation. All had normal or corrected to normal vision. We randomly assigned each participant to one of three conditions (target

### Results

Figure 6 shows a relative frequency histogram of the trials-to-criterion observed across all three conditions. The histogram shows that the majority of participants were able to learn each problem on average in less than 100 trials. However, this histogram also shows that a subset of participants required many more trials to learn each problem. Given that each problem is very simple and that the relevant dimension of each problem is explicitly instructed to participants, it is likely that the participants in the tails of this distribution did not pay attention to the instructions, were not motivated to learn the task, were distracted by other factors, or should be considered outliers for some other reason.

**Figure 6.**
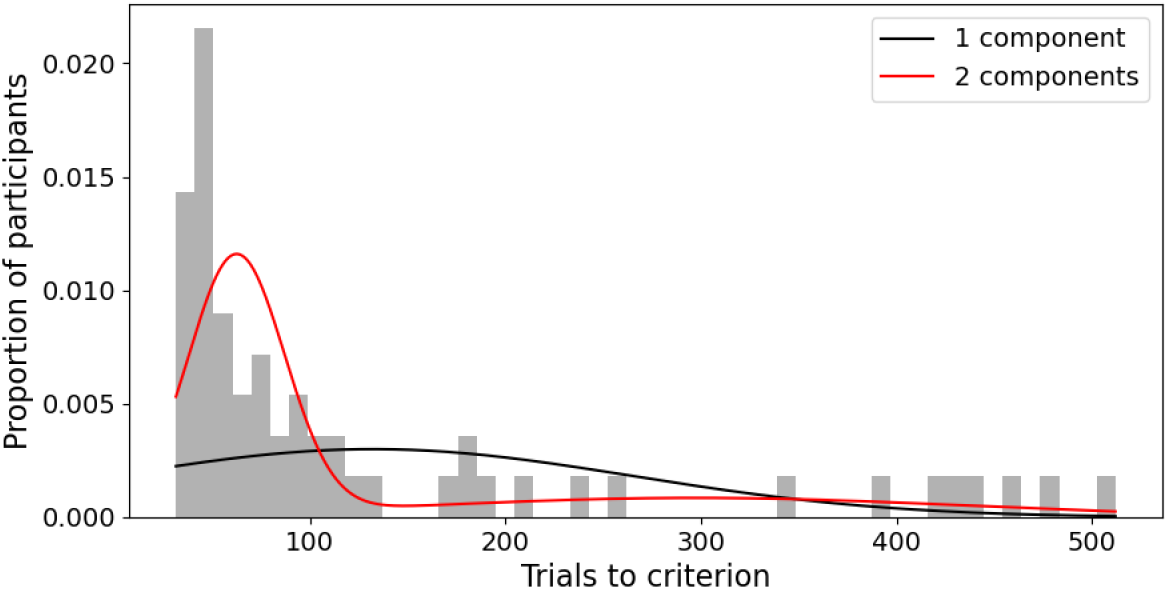
Relative frequency histogram of the trials-to-criterion observed across all three conditions in Experiment 2. The black line are the predictions of the best-fitting one-component Gaussian Mixture Model to these data, whereas the red line represents the best fit of the two-component Gaussian Mixture Model. The two-component model provided a significantly better fit according to AIC, BIC, and a likelihood ratio test.

We modeled the distribution shown in Figure 6 using a Gaussian mixture model with two components. The first component captured participants with a relatively low mean number of trials-to-criterion, while the second component identified outliers according to the above rationale. We compared the fit of this two-component model to a single-component model using the Akaike Information Criterion (AIC) and Bayesian Information Criterion (BIC), both of which indicated a better fit for the two-component model (AIC: 662.81 vs. 735.77; BIC: 673.11 vs. 739.89). Additionally, a likelihood ratio test confirmed that the two-component model provided a significantly better fit than the one-component model (*χ*^2^(3) = 78.96, *p <* .001). Outlier participants were then excluded from further analysis. After making these exlcusions, we were left with the following sample sizes: Control condition (*N* = 11); Delayed-Feedback condition (*N* = 14); Long-ITI condition (*N* = 17).

Figure 7A shows the mean trials-to-criterion in each condition of Experiment 2. A one-way ANOVA revealed a significant effect of condition (*F* (2, 39) = 6.17, *p <* .01, *η*^2^ = .24) and planned comparisons revealed that performance in the Delayed-Feedback condition was significantly worse than in either the Control condition (*t*(23.00) = 2.70, *p <* .05) or the Long-ITI condition (*t*(20.96) = 2.94, *p <* .01). Performance in the Control and Long-ITI conditions did not differ significantly from each other (*t*(17.99) = 0.22, *p* = .83).

**Figure 7.**
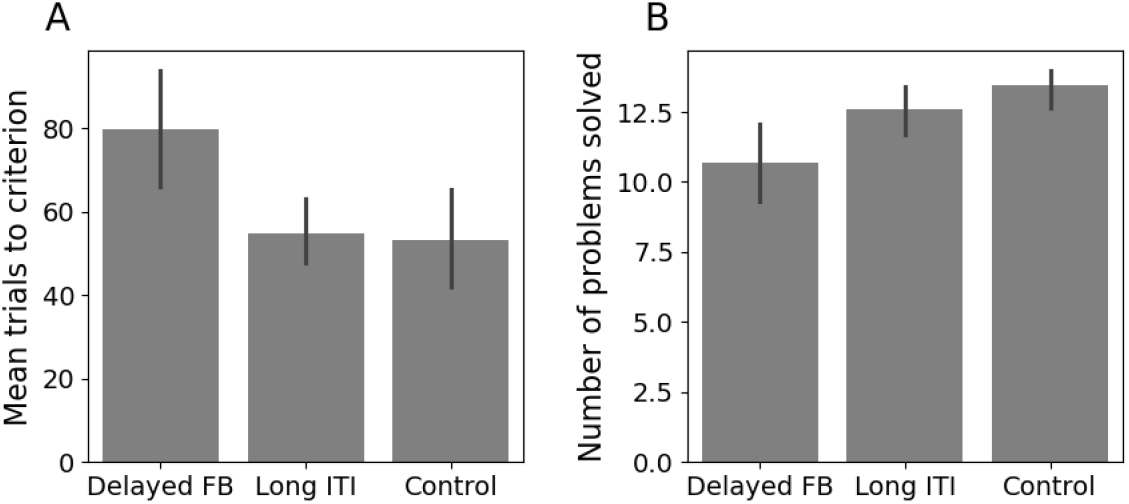
**A**: Mean trials to criterion in each condition of Experiment 2. Error bars are standard errors of the mean. **B**: Mean number of problems solved in each condition of Experiment 2. Error bars are standard errors of the mean.

Figure 7B shows the mean number of problems solved in each condition of Experiment 2. A one-way ANOVA revealed a significant effect of condition (*F* (2, 39) = 5.77, *p <* .01, *η*^2^ = .23). Planned comparisons revealed that performance in the Delayed-Feedback condition was significantly worse than in the Control condition (*t*(18.54) = *−* 3.28, *p <* .01) and worse than in the Long ITI condition (*t*(21.60) = *−* 2.14, *p <* .05). Performance in the Control and Long-ITI conditions did not differ significantly from each other (*t*(25.97) = *−*1.50, *p* = .15).

## Experiment 3

Experiment 3 was designed to reinforce the findings of Experiment 2 by using the same design but with stimuli that have binary-valued dimensions, thereby eliminating criterial learning. Unlike Experiment 2, we did not explicitly instruct participants on the relevant stimulus dimension in Experiment 3. While Experiment 2 isolated criterial learning from rule selection and rule switching, this experiment isolates rule selection and switching from criterial learning. Therefore, Experiment 3 was designed to test whether the impaired learning that we observed in Experiment 2 when the feedback was delayed can be attributed principally to criterial learning or whether it could be due to some more general category-learning process.

### Method

#### Apparatus

The apparatus was the same as in Experiment 2.

#### Stimuli and Categories

The stimuli consisted of colored geometric figures presented on a colored background. These varied across six binary dimensions: the number of items (either one or two), the size of the items (small or large), the color of the items (yellow or blue), the shape of the items (circle or square), the texture of the background (smooth or rough), and the orientation of the background (horizontal or tilted by 20 degrees). This combination resulted in a total of 64 unique stimuli (2^6^). An example trial and stimuli are shown in Figure 8. In all conditions, the order of stimulus presentation was fully randomized for each participant, and the relevant dimension for each problem was selected randomly without replacement from the set of six dimensions.

**Figure 8.**
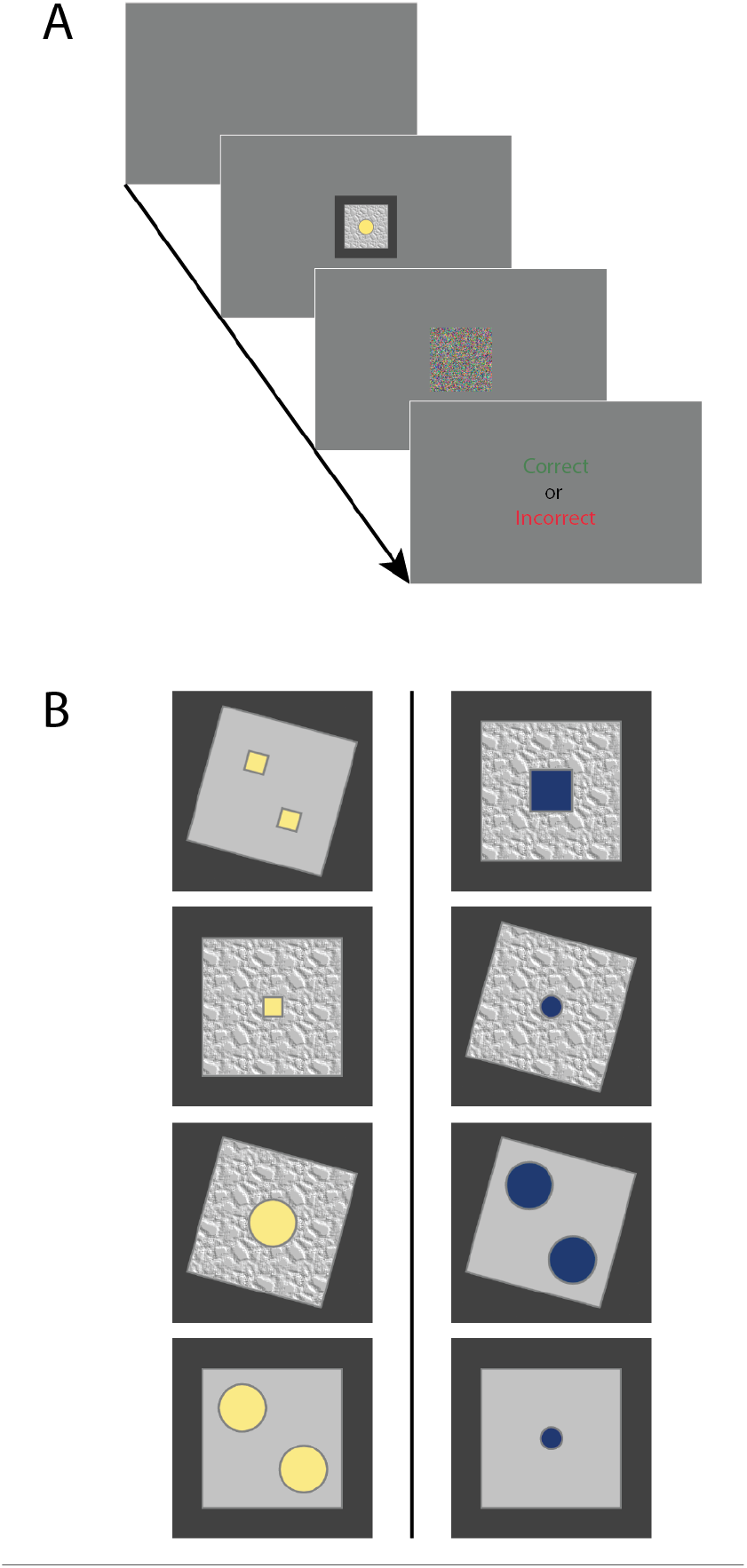
**A:** An example trial from Experiment 3. **B:** Eight example stimuli (out of 64 total stimuli) from a possible one-dimensional rule problem. In this case the correct rule is ‘A’ if the shape color is yellow and ‘B’ if the shape color is blue. The solid line represents the category boundary.

#### Procedure

Participants received detailed instructions about the task. However, unlike in Experiment 2, they were not informed of the relevant dimension or the correct rule. The trial timing for the Delayed-Feedback and Long-ITI conditions in Experiment 3 were the same as in Experiment 2. Experiment 3 did not include a condition that was analogous to the Control condition of Experiment 2. Participants practiced each problem until they responded correctly on 12 consecutive trials. At this point, the problem changed. Each participant completed as many problems as possible in 600 trials or until they had been in the lab for 30 minutes (including time to acquire consent and give instructions), at which point the session was terminated.

#### Participants

Thirty-four participants participated in Experiment 3. All were Macquarie University undergraduates and received course credit for their participation. All had normal or corrected to normal vision. We randomly assigned each participant to one of two conditions: Delayed Feedback (*N* = 17) or Long ITI (*N* = 17).

### Results

Figure 9 shows a relative frequency histogram of the trials-to-criterion observed in both conditions. The histogram shows that the majority of participants were able to learn each problem on average in less than 100 trials. Unlike Experiment 2, there were no highly suspicious outliers in this data set. Nevertheless, for symmetry with Experiment 2, we performed the same Gaussian mixture model analysis as used there. The two-component model provided a slightly smaller AIC value (317.40 vs. 317.78), but a slightly larger BIC value (325.03 vs. 320.83). The likelihood-ratio test was not significant (*χ*^2^(3) = 6.38, *p* = .09). We therefore did not exclude any participants from further analysis in Experiment 2.

**Figure 9.**
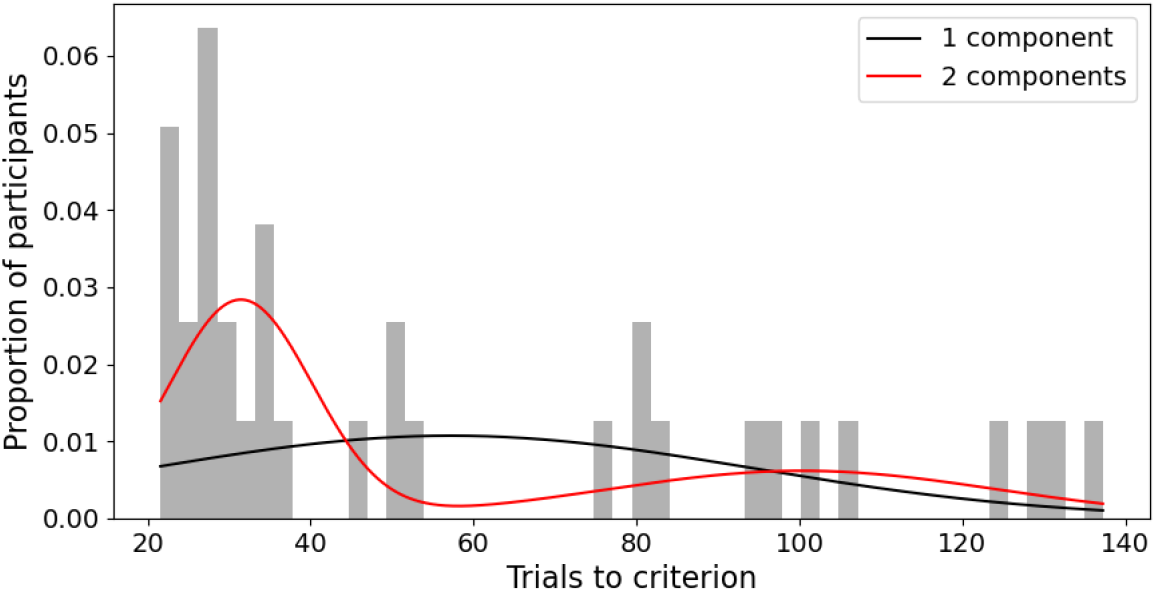
Relative frequency histogram of the trials-to-criterion observed across all three conditions in Experiment 3. The black line represents the best fit of the one-component Gaussian Mixture Model, and the red line represents the best fit of the two-component Gaussian Mixture Model.

Figure 10A shows the mean trials-to-criterion in each condition of Experiment 3. An independent-samples t-test revealed no significant effect of condition (*t*(32.0) = 0.69, *p* = .50, *η*^2^ = .01). The mean number of problems solved by each participant is shown in Figure 10B. An independent-samples t-test revealed no significant effect of condition (*t*(32) = *−*0.49, *p* = .63, *η*^2^ = .01).

**Figure 10.**
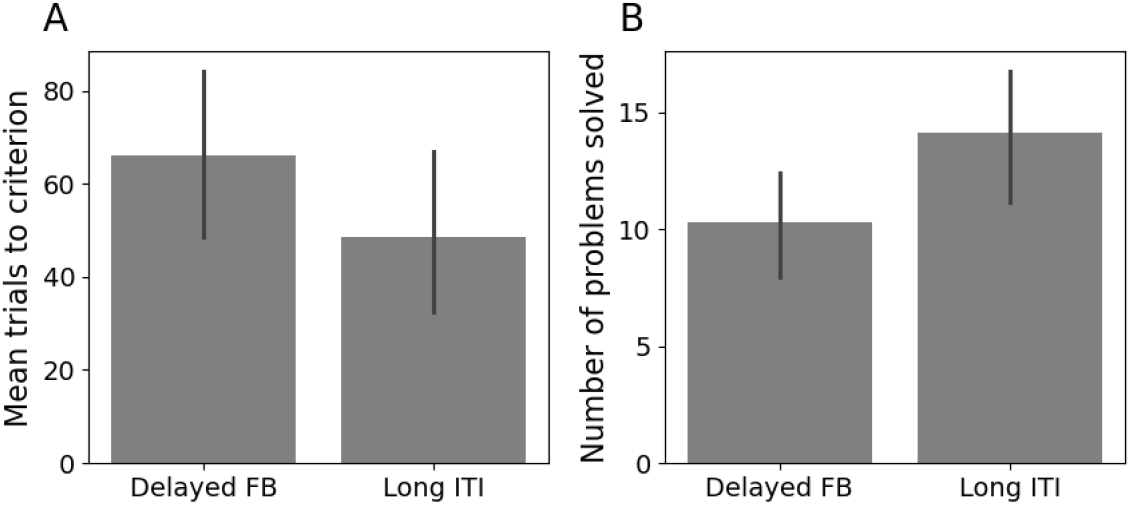
**A**: Mean trials to criterion in each condition of Experiment 3. Error bars are standard errors of the mean. **B**: Mean number of problems solved in each condition of Experiment 3. Error bars are standard errors of the mean.

### Discussion of Experiments 2 and 3

Experiment 2 clearly showed that a short feedback delay of 3.5 s slowed criterial learning. In contrast, increasing the ITI to this same value had no effect on learning. The task we used was one-dimensional category learning, but we isolated criterial learning by instructing participants about the optimal strategy. Specifically, we told them which stimulus dimension was relevant and that there was no trial-by-trial variability on the irrelevant dimension. Although these results strongly suggest that the feedback delay had its interfering effects on criterial learning, Experiment 3 was designed to confirm this inference. The goal here was to examine the effects of the same feedback delays and ITIs on performance in a one-dimensional category-learning task that did not require any criterial learning, but did require category-learning processes that are thought to mediate rule discovery (e.g., rule selection and switching). If the feedback delay effects observed in Experiment 2 were acting on some general category-learning skill then we should have seen the same interfering effects of feedback delay in Experiment 3. However, if the feedback-delay interference of Experiment 2 was operating selectively on criterial learning, then it should disappear in Experiment 3, since no criterial learning was required. Our results strongly supported this latter prediction. Therefore, Experiments 2 and 3 together strongly suggest that criterial learning is impaired by feedback delays and is relatively unaffected by the length of the ITI.

Two prior studies investigated similar issues. First, Ell and colleagues reported that feedback delay and also a concurrent memory-scanning task each impaired rule-based category learning (Ell et al., 2009). Based on these results, they hypothesized that criterial learning may be tied to working-memory capacity and therefore to explicit cognitive mechanisms. However, criterial learning was confounded with rule selection and task difficulty in their design. Furthermore, Ell et al. (2009) found that feedback delay only impaired learning when working memory demand was high – that is, when participants had to learn more than one response criterion for optimal performance. In contrast, we found that feedback delay impairs learning even when working memory demands are trivial (participants never have to keep in mind more than one criterion).

Second, Bohil and Wismer (2014) reported that the effects of unequal base rates on criterion placement in rule-based category learning were diminished under delayed feedback and also under an observational training protocol that is also thought to selectively impair basal ganglia-dependent associative-learning mechanisms. These results are consistent with our findings. However, because they were obtained by manipulating base rates, it is difficult to conclude that the delay affected criterial learning, *per se*. For example, consider a simple two-stage model in which the first stage learns the base rates and the second stage uses what the first stage learned to adjust the response criterion appropriately. The Bohil and Wismer (2014) results are also consistent with the hypothesis that the feedback delay impaired the first of these stages but not the second. Our results strongly suggest that feedback delays impair criterial learning (and therefore the second of these hypothetical stages).

## General Discussion

We developed three novel models of criterial learning that were designed to explore how feedback delay and ITI duration affect criterial learning. Two of these models assumed that decisions are made by comparing the current percept to a stored referent, or criterion, and that the remembered value of this criterion is updated following error feedback via a gradient-descent learning rule. The first of these models, the time-dependent drift model, posits that both the criterion and the percept drift over time, leading to similar impairments when feedback is delayed or the ITI is increased. The second model, the delay-sensitive learning model, assumes that although neither the percept nor the criterion drift over time, optimal criterial learning requires immediate feedback. As a result, this model predicts that increasing the feedback delay will slow the rate of criterial learning. The third model, the reinforcement-learning model, assumes that criterial learning arises through the gradual formation of stimulus-response associations, rather than via the construction of a response criterion. This model also assumes that the rate at which these associations are learned is slowed by feedback delays.

Simulations of these models in a task that was structurally identical to Experiment 2 revealed that the time-dependent drift model typically predicts that the most important variable for criterial learning is the total amount of time between the response and the stimulus presentation that defines the next trial. Where the feedback is presented within this interval is relatively unimportant. In contrast, both the delay-sensitive learning model and the reinforcement-learning model consistently predict that feedback delays should impair performance more than a long ITI.

The experiments were designed to test these predictions. Our results strongly suggested that human criterial learning is sensitive to feedback delay but not to the ITI duration. This suggests that the delay-sensitive learning model and the reinforcement-learning model provide a better account of human behavior across wide ranges of their respective parameter spaces than the time-dependent drift model.

The time-dependent drift model was based on the idea that criterial learning might be entirely supported by working memory. The assumption of drift in this model stems from the understanding that maintaining items in working memory is inherently challenging. The longer an item is held, the more likely it is to deteriorate. Therefore, the model predicts that performance should deteriorate as the time between the response on trial *n* and stimulus presentation on trial *n* + 1 increases, regardless of how this interval is divided between feedback delay and ITI. The results of Experiments 1 and 2 strongly disconfirm this prediction.

In contrast, the delay-sensitive learning model was inspired by the idea that a response criterion might be encoded in a more stable memory system. One appealing candidate for this function is the cerebellum, where synaptic plasticity has been shown to follow a gradient descent learning rule and also to be sensitive to feedback timing (Brudner et al., 2016; Held et al., 1966; Honda et al., 2012; Kitazawa et al., 1995; Kitazawa & Yin, 2002). Finally, the reinforcement-learning model draws on the idea that criterial learning could emerge through stimulus-response association learning driven by dopamine-dependent synaptic plasticity in the striatum. This process aligns with established theories of category learning that posit a key role to this learning process in the basal ganglia Ashby et al., 1998.

Our results do not strongly favor the delay-sensitive learning model over the reinforcement learning model or vice versa. The critical disagreement between these accounts is whether the criterion has any real psychological meaning. In the delay-sensitive learning model it does, and thus this model is psychologically similar to the time-dependent drift model – the main disagreement being about whether the representation of the criterion is stable and if the updating of the criterion is sensitive to feedback delay.

The reinforcement-learning model makes very different psychological assumptions. This model assumes that behaviors described as “criterial learning” are actually mediated by the learning of stimulus-response associations. According to this account, the criterion has no internal representation and therefore no psychological meaning. Testing between these two very different accounts of criterial learning should be a focus of future research.

We are not aware of any data that directly addresses the question of whether a response criterion is a fundamental psychological construct, as the delay-sensitive learning model suggests, or whether it is unnecessary, as assumed by the reinforcement learning model. Even so, there are some results in the category-learning literature that support the interpretation of the reinforcement-learning model. In information-integration (II) categorylearning tasks, the optimal strategy is similarity-based, and difficult or impossible to describe verbally (e.g., Ashby & Valentin, 2018). When the stimuli vary on two dimensions, the stimuli from contrasting categories can be partitioned by a decision bound that is conceptually similar to a response criterion. In both cases, all stimuli on one side are associated with one response, and all stimuli on the other side are associated with the contrasting response. Furthermore, in agreement with Experiment 2, II category learning is impaired by short feedback delays (Maddox & Ing, 2005; Maddox et al., 2003). The analogous question in II category learning is whether the decision bound is learned directly or whether it is simply the set of points that divide the perceptual space into contrasting response regions. In fact, the evidence strongly supports this latter interpretation (Ashby & Waldron, 1999; Casale et al., 2012). For example, if the decision bound is learned, then it should be possible to apply this bound to novel stimuli. With rule-based categories, this is easy for participants, but with II categories there is no evidence that the response strategy that participants learn can be generalized to novel stimuli (Casale et al., 2012) – a result that strongly supports the hypothesis that the decision bound has no psychological meaning.

Our results clearly demonstrate that criterial learning is impaired by delayed feedback, and not by extending the intertrial interval. These results are consistent with the hypothesis that criterial learning is a form of basal-ganglia mediated associative learning, and are inconsistent with hypotheses that criterial learning is a working-memory-based process. Thus, our results provide a critical constraint on future models of rule-based classification and decision making, and possibly also on more general accounts of criterion setting, such as in signal detection theory.

Previous studies have failed to find that feedback delays impair rule-based category learning, and on the face of it, this seems to contradict our finding that feedback delays impair criterial learning. However, all earlier RB studies that looked for effects of a feedback delay, either used binary-valued stimulus dimension and so no criterial learning was needed, or else set the response criterion exactly midway between the category prototypes, which makes criterial learning trivial. For example, under these conditions, criterial learning might not even require feedback. The unsupervised category-learning experiments reported by Ashby et al. (1999) provide strong support for this because all of their rule-based participants learned the correct criterion (which was midway between the category means), even though the task was completely unsupervised. Moreover, all previous studies failed to isolate criterial learning, so even if there was an effect of feedback delay on criterial learning, it could have been masked by larger effects caused by other rule-learning processes.

Criterial learning is among the most classic and ubiquitous of all cognitive skills. For example, signal-detection theory teaches that it is the central form of learning in an enormous range of decision-making tasks – everything from simple YES-NO detection of a weak signal to assessing the guilt or innocence of a defendant in a jury trial. Our results suggest that even in simple rule-based tasks, criterial learning seems to be subserved, at least in part, by associative mechanisms. Most current theories tend to classify tasks as either executive function (e.g., rule-based category learning) or procedural (e.g., mirror tracing). Our results suggest that such classification schemes might oversimplify how humans perform these tasks, and therefore that much more work is needed to understand how different learning and memory systems interact.

## Transparency and Openness

All data have been made publicly available in the following GitHub repository: https://github.com/crossley/crit_learn_delay

## Author Notes

Preparation of this article was supported by Public Health Service Grant MH2R01063760.

## Notes

https://github.com/crossley/crit_learn_delay

### Competing Interest Statement

The authors have declared no competing interest.

